# Quantitative Image Analysis of Tissue Properties: A MATLAB Tool for Measuring Morphology and Co-localization in 2D Images

**DOI:** 10.1101/2024.04.03.587971

**Authors:** Shira Landau, Erez Shor, Milica Radisic, Shulamit Levenberg

## Abstract

In recent years, the structural analysis of tissue elements has gained significant importance in biomedical research. Advancements in imaging technologies have created a pressing need to quantify tissue and cell properties accurately. This paper introduces a MATLAB-based analytical tool designed to measure a spectrum of properties from 2D images of tissues and cells. Our software efficiently computes parameters such as eccentricity, orientation, density, co-localization, size, and perimeter. The algorithm’s precision in evaluating these characteristics has broad implications for enhancing the understanding of various biological processes and diseases. The code’s flexibility allows for application across different tissue types and experimental conditions, providing researchers with a robust method for quantitative analysis. This advancement in computational image analysis represents a pivotal step towards more detailed and objective assessment in tissue engineering and cellular biology.

## Introduction

The quantification of structural properties of tissue elements has emerged as a pivotal aspect of modern biomedical research, significantly aiding in the understanding of complex biological systems and disease pathologies. As imaging technology has advanced, the need for precise measurement of these properties has grown, necessitating the development of sophisticated computational tools. In this study, we introduce a MATLAB code capable of quantifying key properties of tissue or cell elements from 2D images.

Quantitative image analysis has become indispensable in tissue engineering, developmental biology, and pathology. It allows researchers to characterize cellular morphology, identify phenotypic changes in response to treatments, and understand tissue architecture and its functional implications^1-3^. Traditional methods of analysis, often manual or semi-automated, are not only time-consuming but also prone to subjective bias, underscoring the necessity for automated, reliable, and reproducible computational techniques^1^.

The MATLAB code presented in this paper addresses these challenges by automating the measurement of eccentricity, which quantifies the deviation of a cell’s shape from a perfect circle^4^; orientation, which determines the angle of the major axis of the cell’s ellipse to a reference axis; density, which assesses the number of cells per unit area; co-localization, which evaluates the spatial overlap of different cellular markers; size, which measures the area covered by a cell or group of cells; and perimeter, which calculates the boundary length of the cells.

This comprehensive approach to analyzing tissue and cell images leverages MATLAB’s robust computational capabilities, allowing for the processing of large datasets with precision and efficiency. The code is designed to be user-friendly, requiring minimal user input and providing clear, interpretable results that can be further analyzed statistically. It is adaptable to a wide range of tissues and cells, making it an invaluable tool for multiple research domains.

Moreover, this tool’s introduction into the scientific community has the potential to standardize quantitative analysis methodologies across laboratories, fostering consistent and comparative studies. By automating complex analysis, it enables researchers to focus on experimental design and interpretation of results, ultimately accelerating the pace of discovery.

In summary, the MATLAB code developed for this study stands as a significant contribution to the field of quantitative image analysis. Its implementation will facilitate a deeper understanding of the structural properties of tissues and cells, paving the way for breakthroughs in biomedical research and diagnostic procedures.

### Procedure

#### Image Collection

In this study, the MATLAB code was developed to analyze 2D images of tissues and cells of RGB images. These images can be either JPEG or TIFF formats at any resolution or magnification. When comparing between groups or treatments, images should be captured using uniform acquisition parameters, such as laser gain and exposure settings, as well as maintaining consistent magnification and image size. The use of a scale bar should be avoided. If images exhibit a biased signal due to different staining protocols or tissue thickness, it could skew the results. This issue can be addressed by applying contrast enhancement to the dimmer images.

#### Matlab requirement

Any version of MATLAB from 2013 onwards can be used, and the Image Processing Toolbox should also be installed.

#### MATLAB Code Description

1. Reading the image

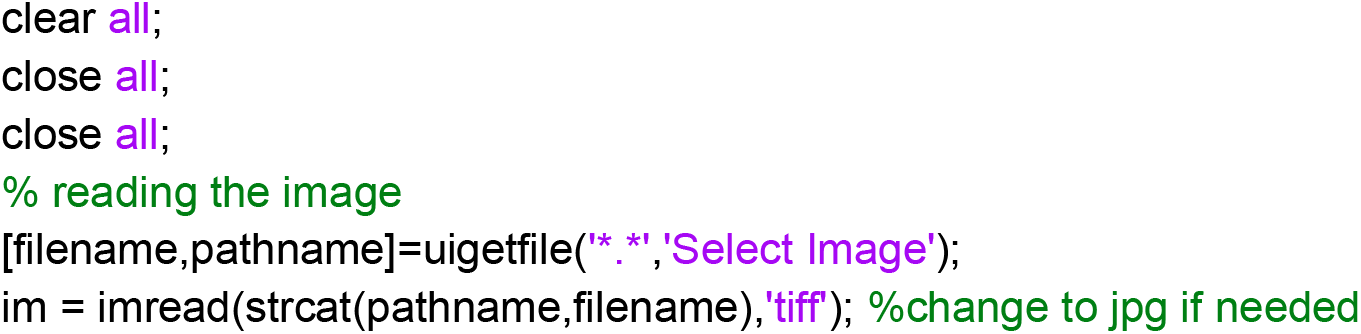

2. Selecting a region of interest (ROI):

If only part of the image is analyzed and a manual selection is required, this part of the code would be used:

~~~
         ROI = roipoly(im); %for manual selection use this
~~~

If the whole image is being analyzed

~~~
         ROI=1;
~~~

3. Dividing the image into 3 RGB channels:

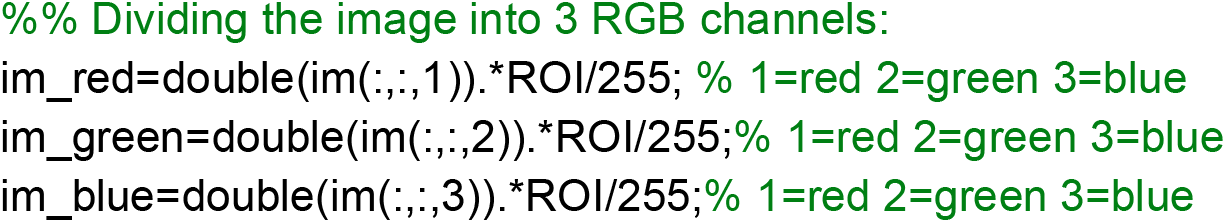

4. Converting the images into binary images using a defined threshold

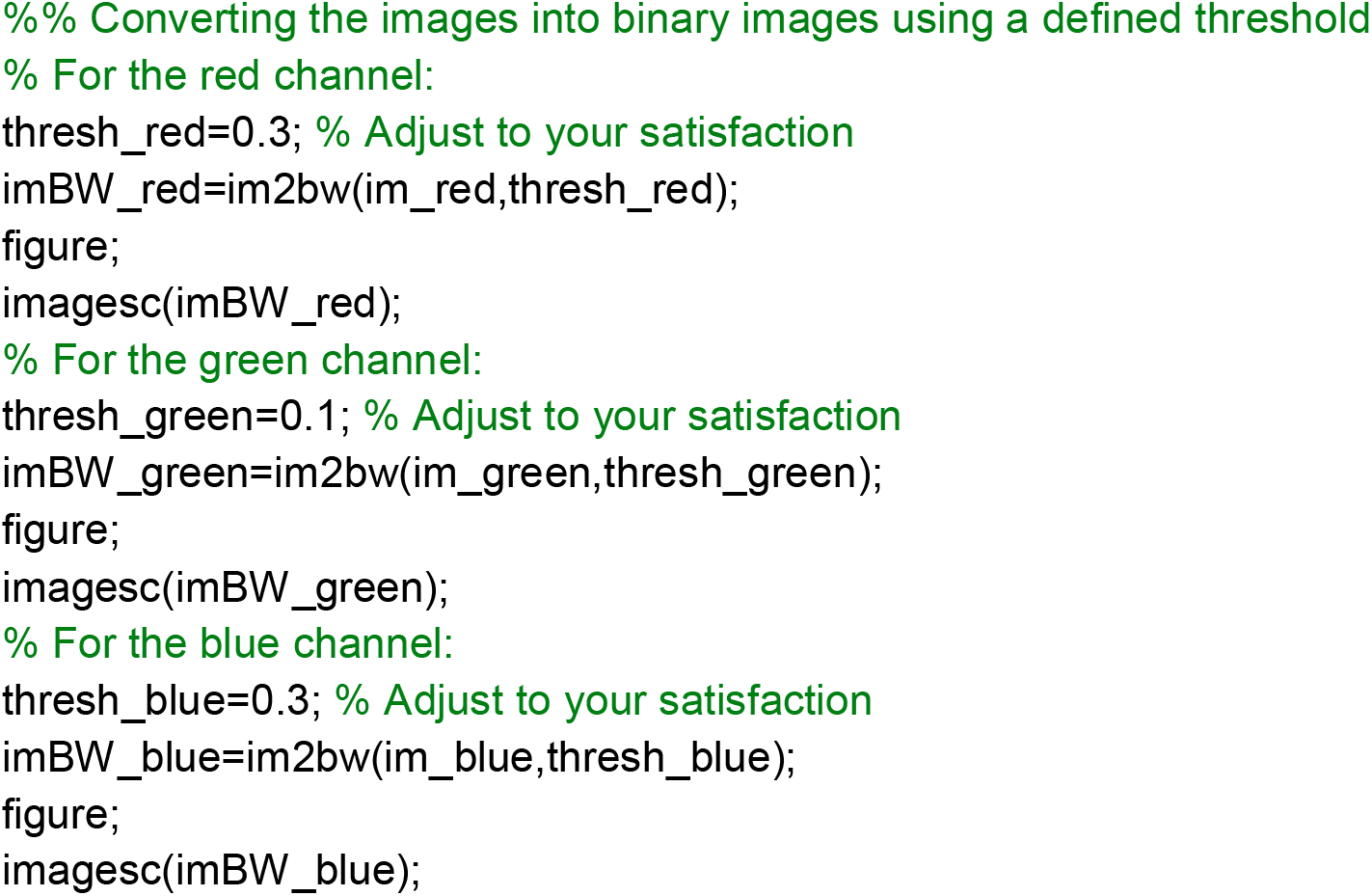

For co-localization analysis:

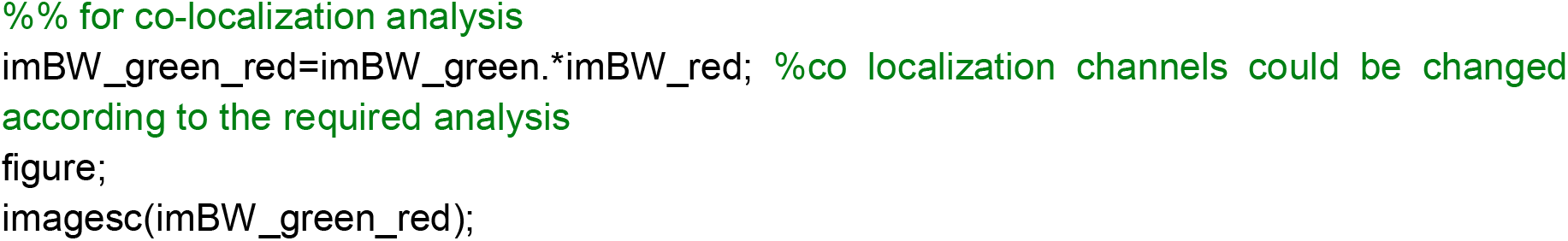

Note: Images are displayed at this point for you to fine-tune the thresholding parameters until satisfied.

1. Measure Image properties with the regioprops function:

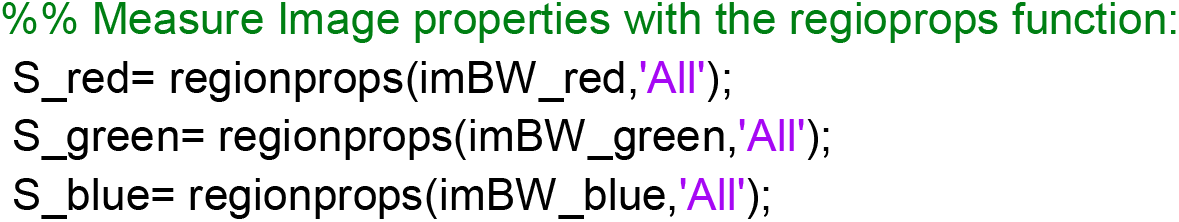
2. Image properties of the red channel

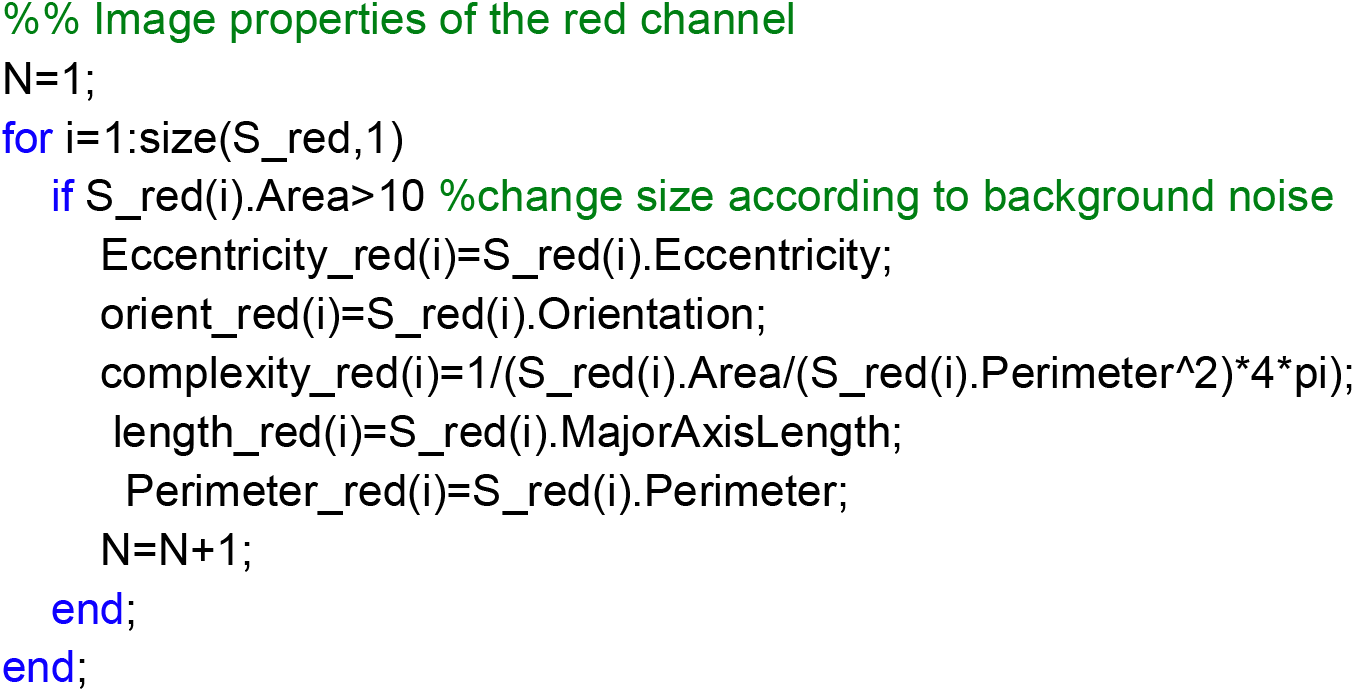
3. Image properties of the green channel

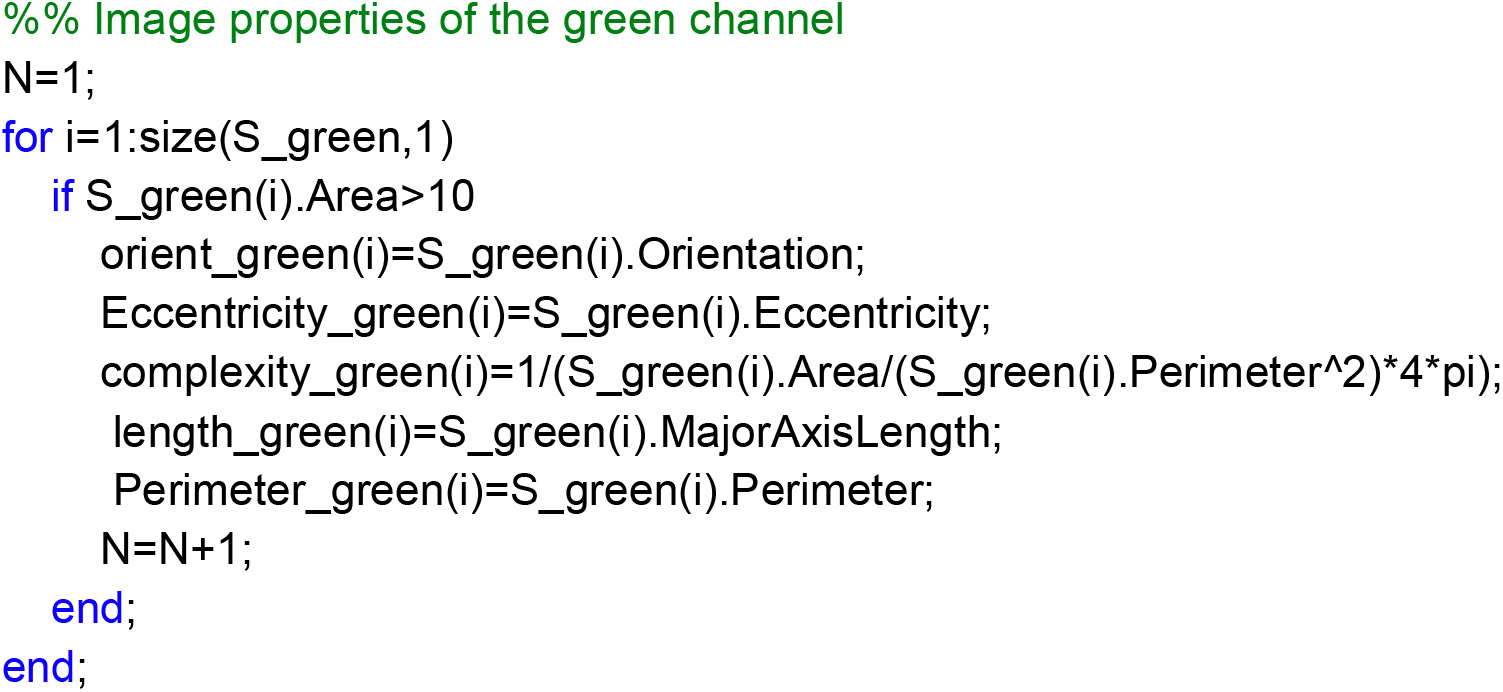
4. Image properties of the blue channel

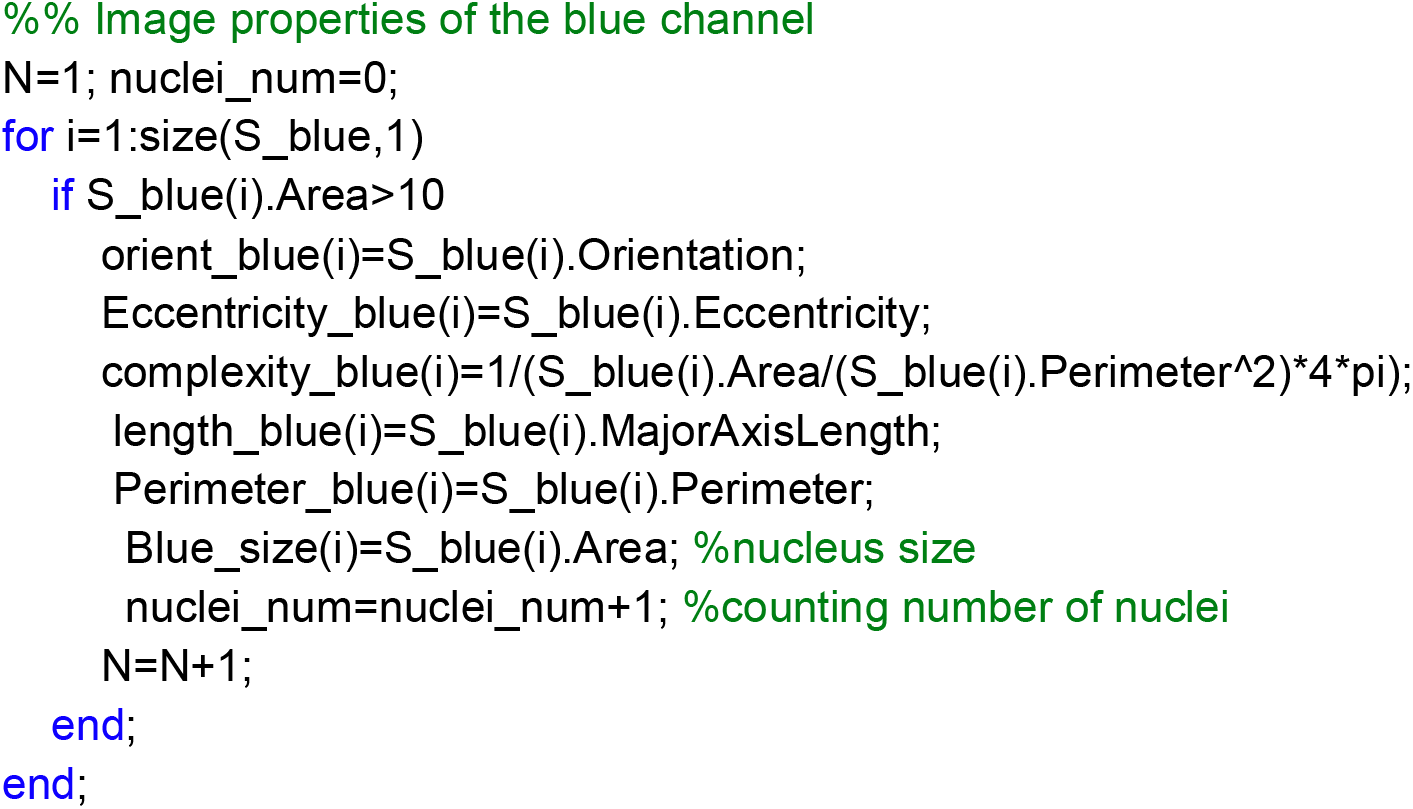 Note 1: change the area size filter according to your noise particle size.
5. Averages calculation for all parameters and channels

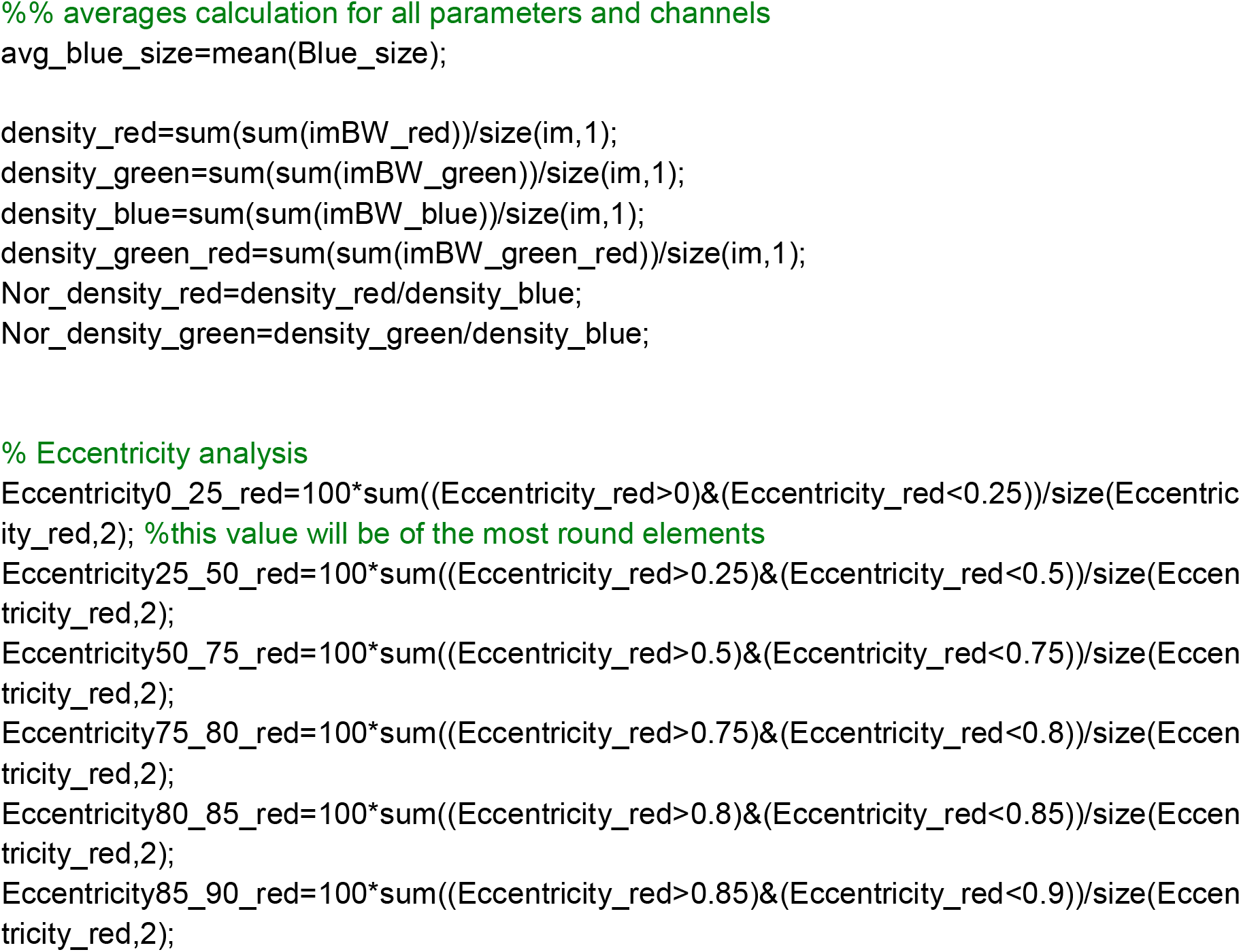

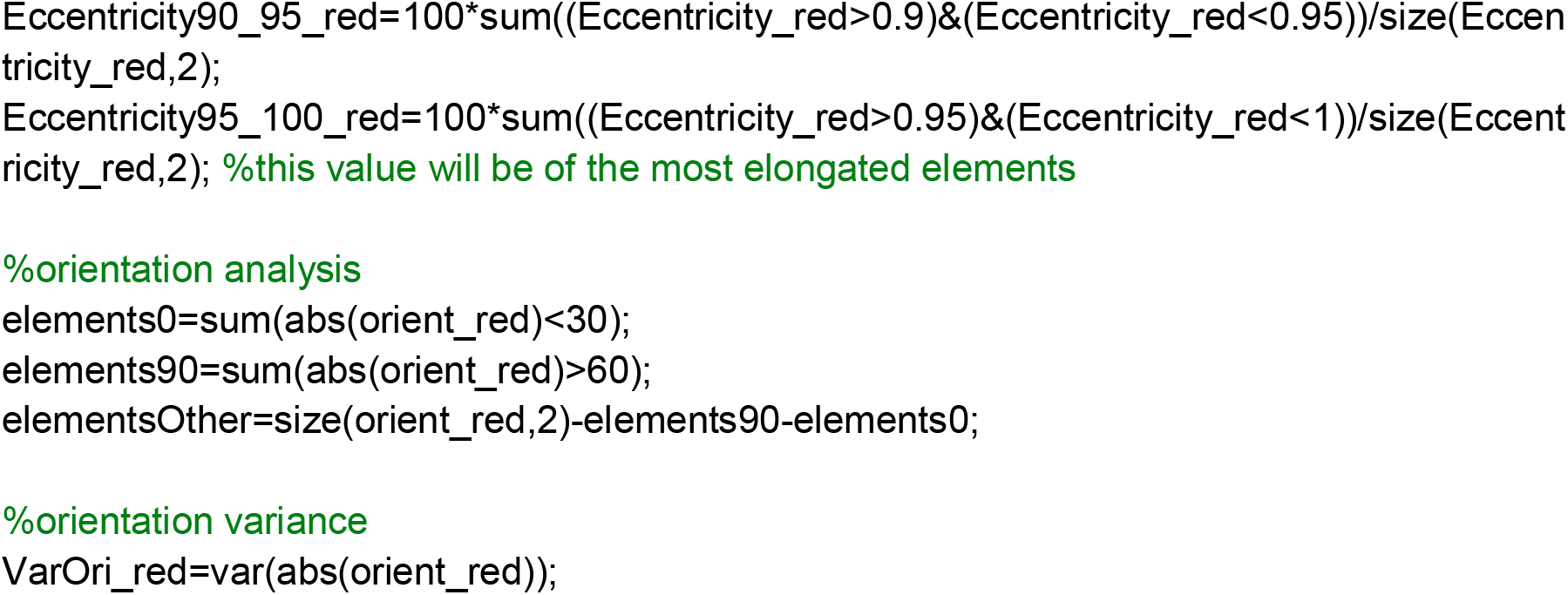
6. Writing the results into a CSV file:

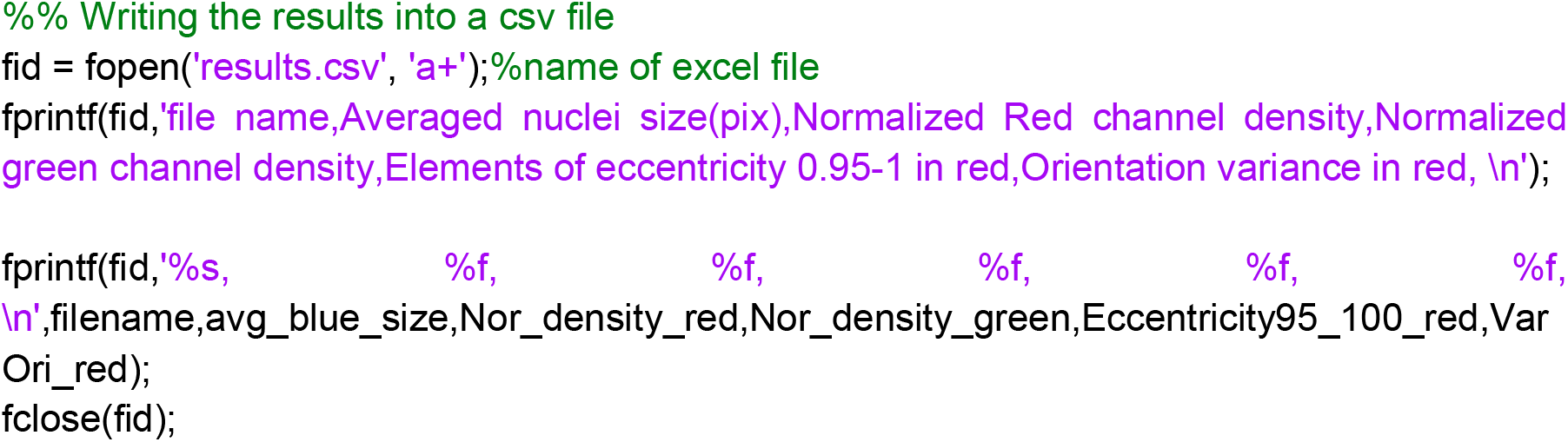

#### If multiple images are used, the “*Image Batch Processor*” APP is recommended. Using this adjusted function

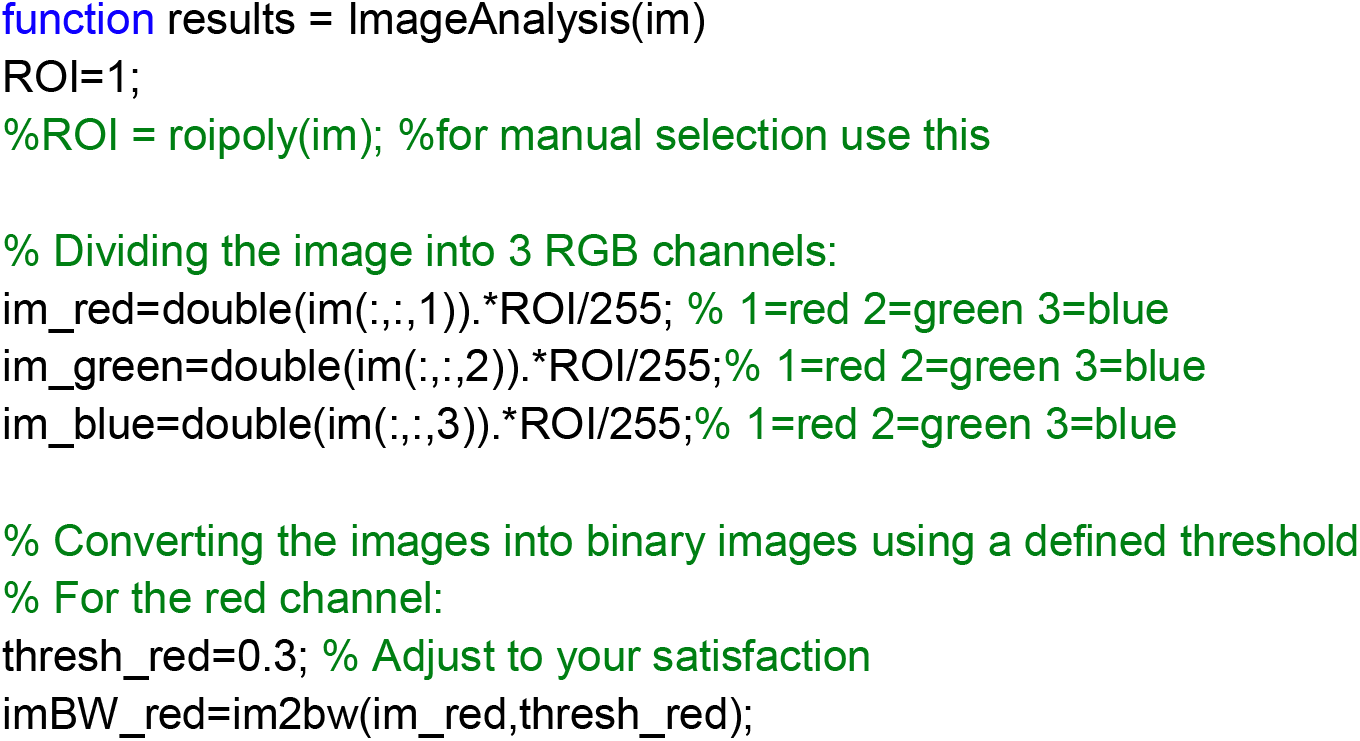

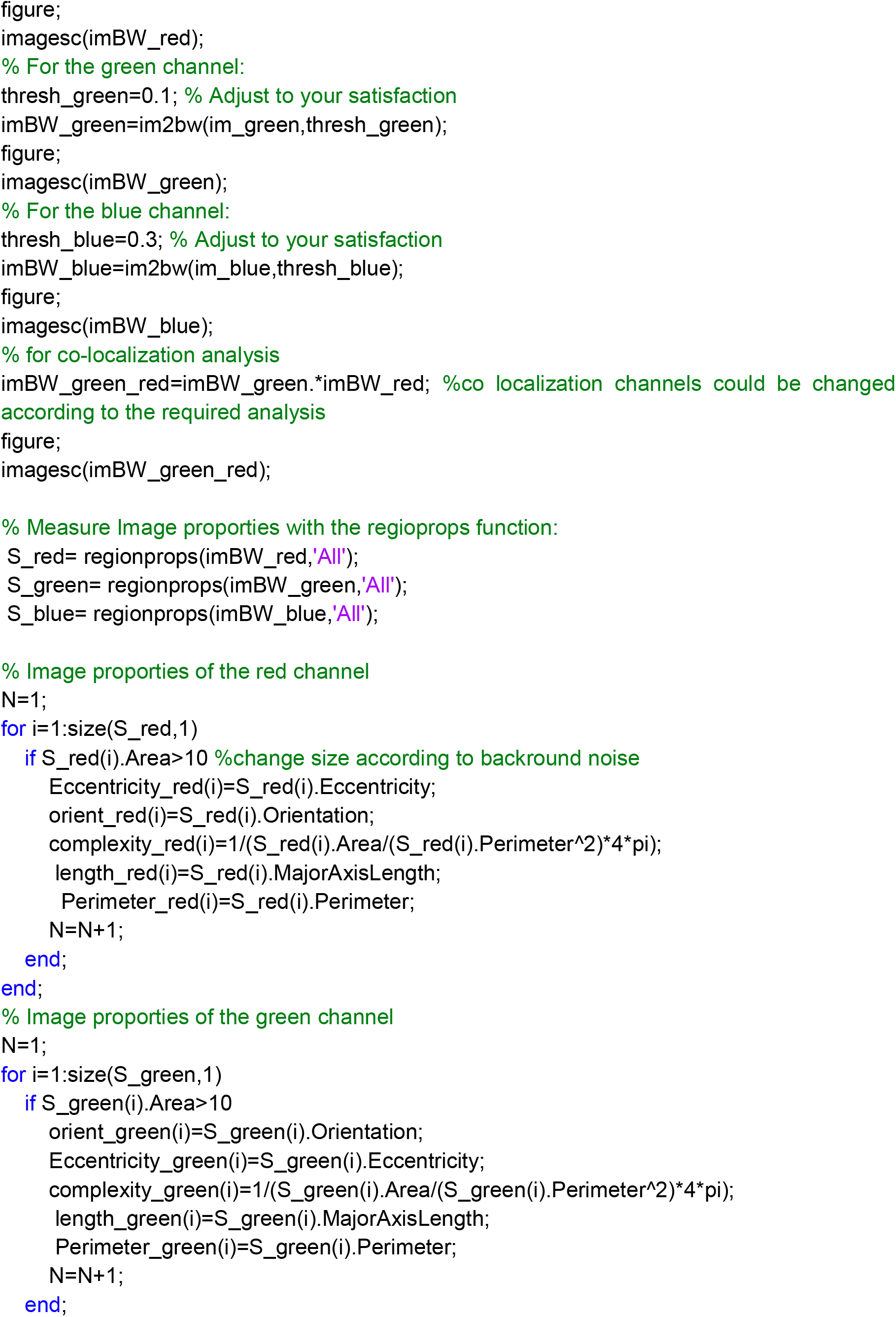

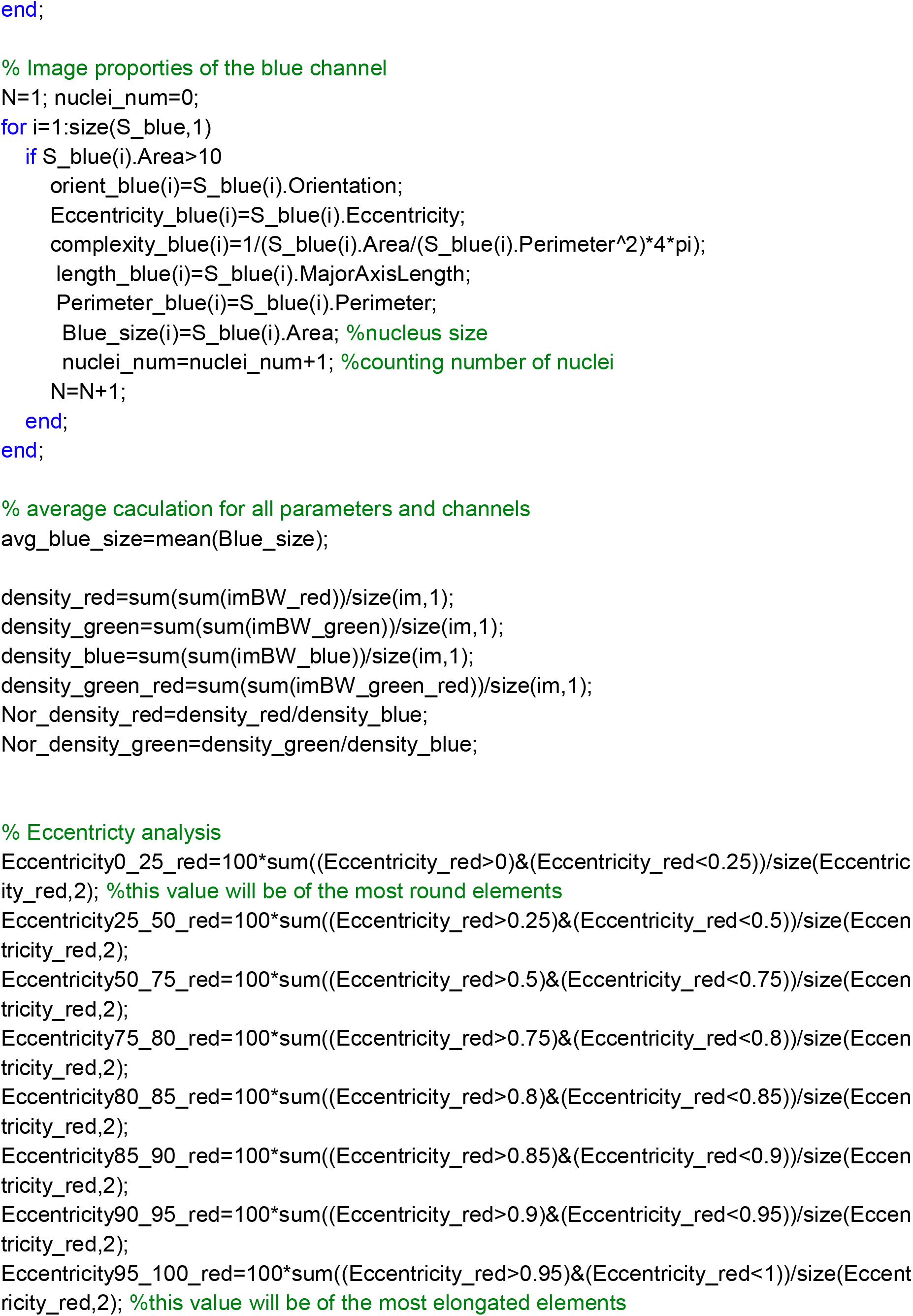

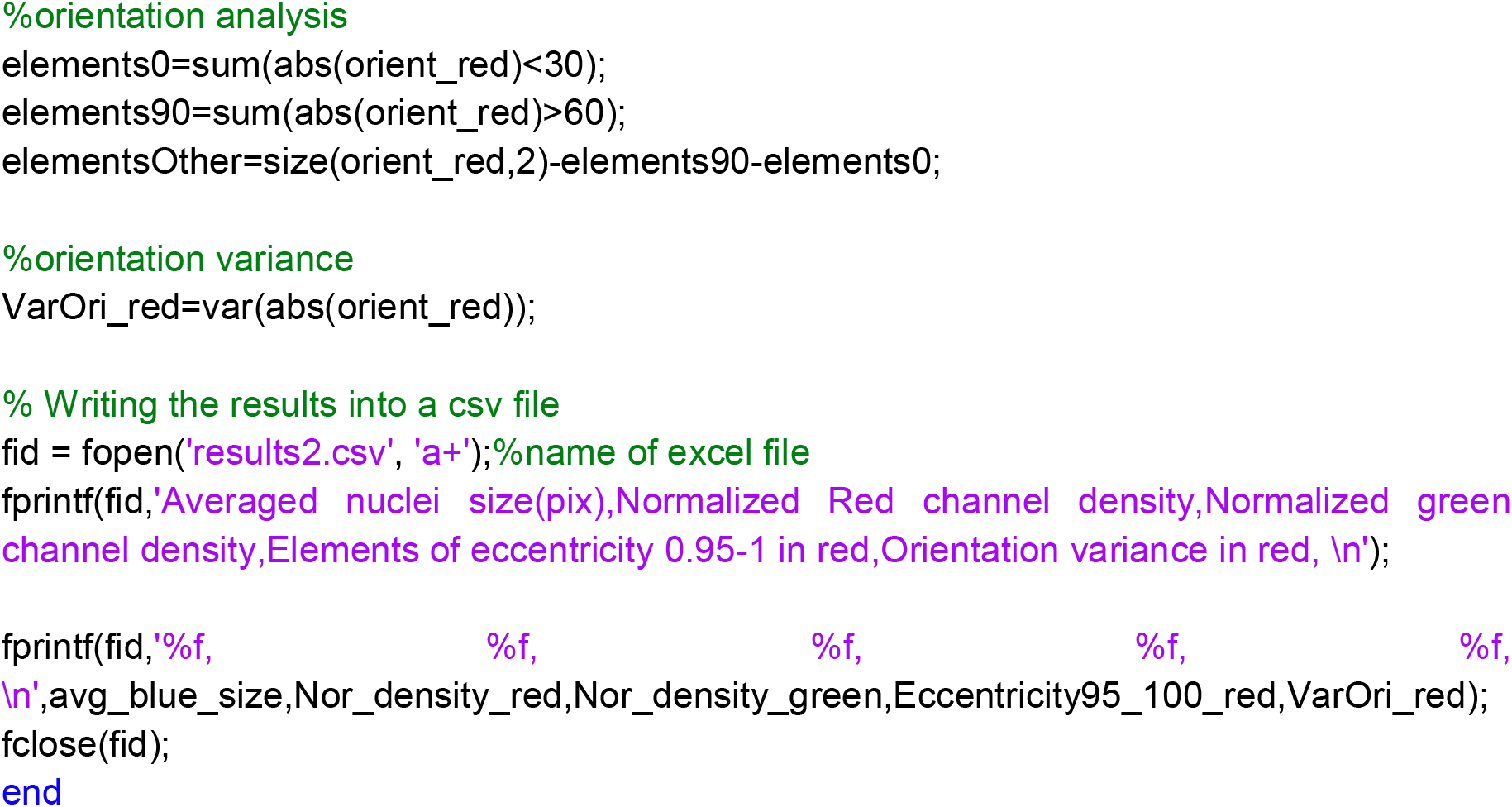

### Example

This example outlines the measurement of elements within an in vitro tissue composed of GFP-expressing endothelial cells, dental pulp stem cells (DPSCs), and cardiomyocytes embedded in a gel and stained with the cardiac marker MLC2V.

Subsequently, the image is segmented into three binary images representing each fluorescent channel (**Figure 1**).

**Figure 1:**
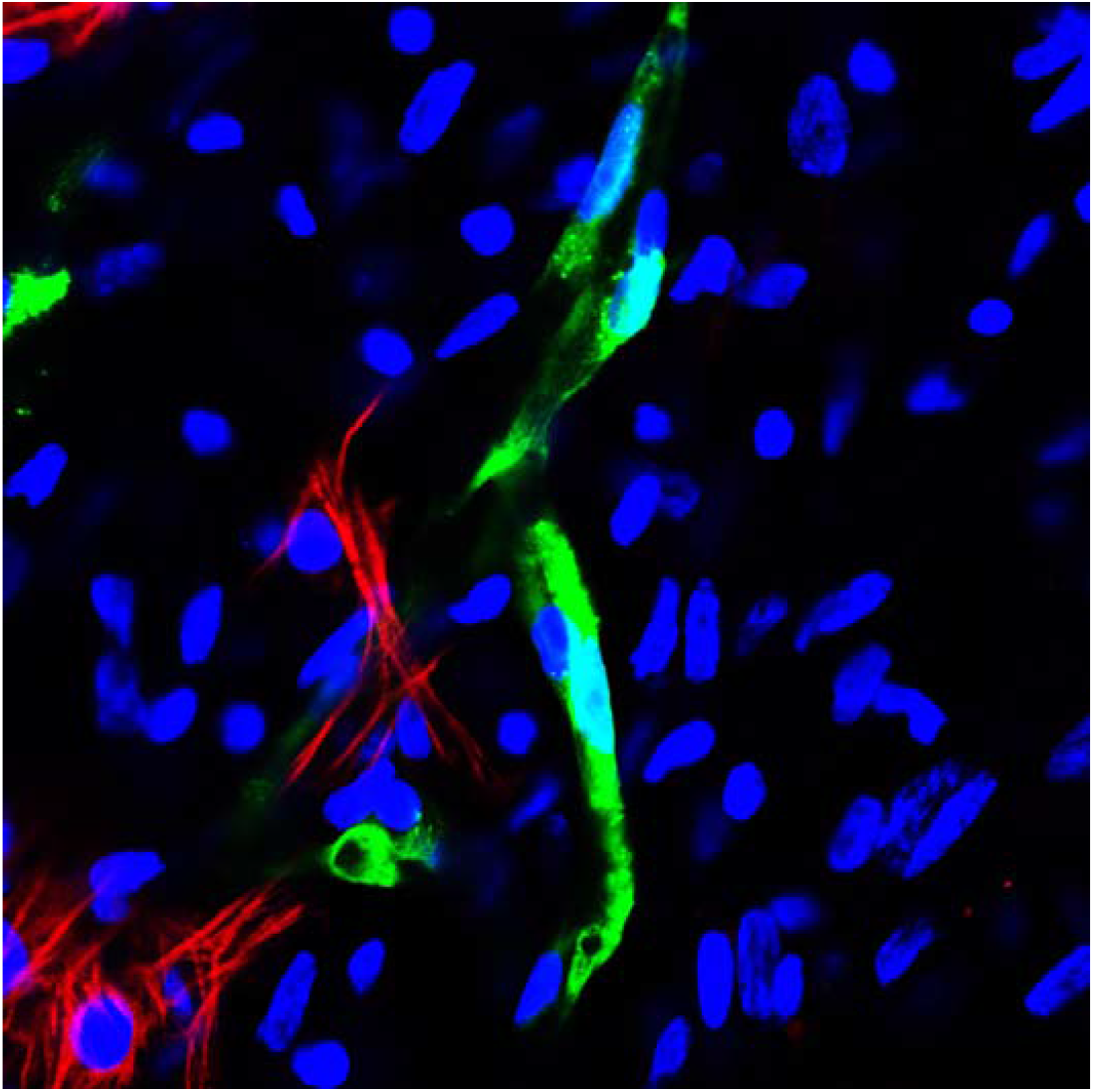
in vitro culture featuring GFP-expressing endothelial cells (green), cardiomyocytes, and DPSCs, stained with DAPI (blue) and MLC2V (red).

These images are processed using the *regionprops* function to extract the necessary measurements, which are then cataloged in an Excel spreadsheet, organizing each measurement into separate columns and each sample into distinct rows (**Figure 2**).

**Figure 2:**
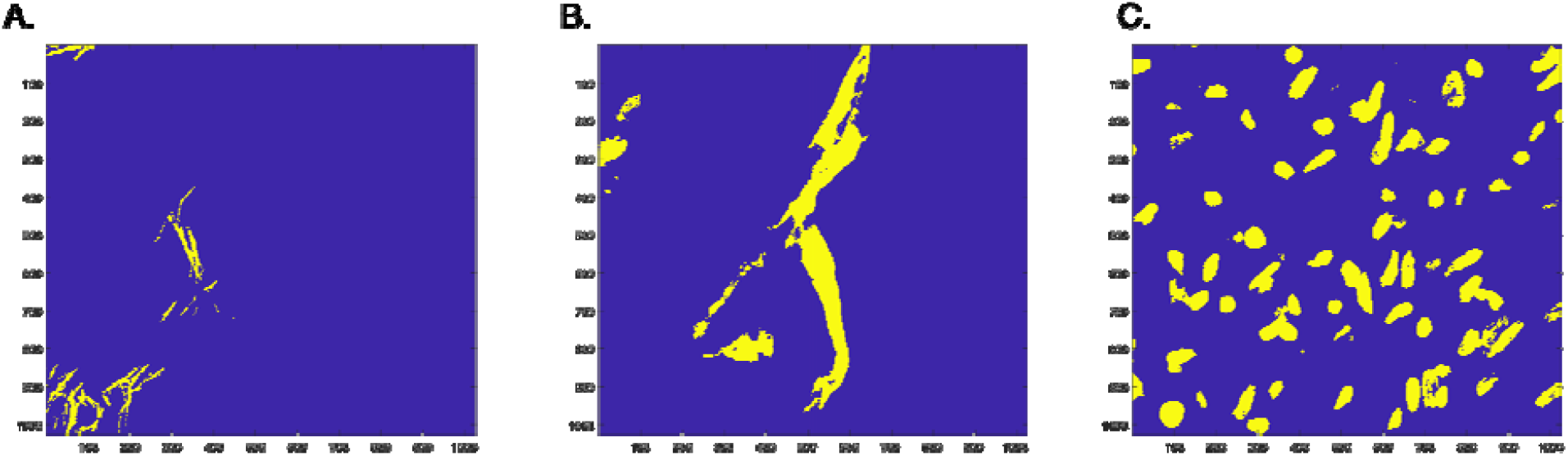
Binary images of the three channels following appropriate thresholding: **A**. red, **B**. green, and **C**. blue channels.

**Figure 3:**
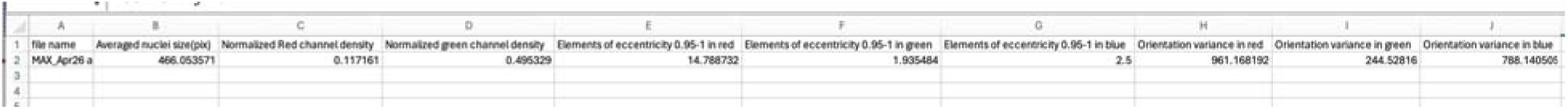
the results table from the image analysis. In this example, the following parameters were measured: average nucleus size, red channel density normalized to nuclei, green channel densit normalized to nuclei, the percentage of elements with an eccentricity value between 0.95 and 1 in the red, green, and blue channels, and the variance in orientation across the red, green, and blue channels.

The results indicate that the normalized density of the green channel exceeds that of the red channel, as expected. Additionally, the red channel, as anticipated, contained the highest percentage of elements with an eccentricity value between 0.95 and 1. In terms of orientation, the green channel exhibited the lowest variance, indicating the most aligned features (F**igure 3**).

### Summary

The study introduces a MATLAB-based analytical tool capable of accurately measuring a range of properties from 2D images of tissues and cells, such as eccentricity, orientation, density, and size. Enhanced by a batch processing feature, the software can swiftly analyze hundreds of images, offering a significant increase in efficiency and making it a robust and versatile solution for comprehensive quantitative analysis in tissue engineering and cellular biology.

## Supporting information

Code and function

