## Supplementary material for "Quantitative Image Analysis of Tissue Properties: A MATLAB Tool for Measuring Morphology and Co-localization in 2D Images": Code and function: code.m

```
clear all;
close all;
close all;
% reading the image 
[filename,pathname]=uigetfile('*.*','Select Image');
im = imread(strcat(pathname,filename),'tiff'); %change to jpg if needed

%% Selecting a region of interest (ROI):
ROI=1;     
%ROI = roipoly(im); %for manual selection use this

%% Dividing the image into 3 RGB channels:
im_red=double(im(:,:,1)).*ROI/255; % 1=red 2=green 3=blue
im_green=double(im(:,:,2)).*ROI/255;% 1=red 2=green 3=blue
im_blue=double(im(:,:,3)).*ROI/255;% 1=red 2=green 3=blue

%% Converting the images into binary images using a defined threshold
% For the red channel:
thresh_red=0.3; % Adjust to your satisfaction
imBW_red=im2bw(im_red,thresh_red);
figure;
imagesc(imBW_red);
% For the green channel:
thresh_green=0.1; % Adjust to your satisfaction
imBW_green=im2bw(im_green,thresh_green);
figure;
imagesc(imBW_green);
% For the blue channel:
thresh_blue=0.3; % Adjust to your satisfaction
imBW_blue=im2bw(im_blue,thresh_blue);
figure;
imagesc(imBW_blue);
%% for co-localization analysis
imBW_green_red=imBW_green.*imBW_red; %co localization channels could be changed according to the required analysis
figure;
imagesc(imBW_green_red);

%% Measure Image proporties with the regioprops function: 
 S_red= regionprops(imBW_red,'All');
 S_green= regionprops(imBW_green,'All');
 S_blue= regionprops(imBW_blue,'All');

%% Image proporties of the red channel
N=1;
for i=1:size(S_red,1)
    if S_red(i).Area>10 %change size according to backround noise
        Eccentricity_red(i)=S_red(i).Eccentricity;
        orient_red(i)=S_red(i).Orientation;
        complexity_red(i)=1/(S_red(i).Area/(S_red(i).Perimeter^2)*4*pi);
         length_red(i)=S_red(i).MajorAxisLength;
          Perimeter_red(i)=S_red(i).Perimeter;
        N=N+1;
    end;  
end;
%% Image proporties of the green channel
N=1;
for i=1:size(S_green,1)
    if S_green(i).Area>10
        orient_green(i)=S_green(i).Orientation;
        Eccentricity_green(i)=S_green(i).Eccentricity;
        complexity_green(i)=1/(S_green(i).Area/(S_green(i).Perimeter^2)*4*pi);
         length_green(i)=S_green(i).MajorAxisLength;
         Perimeter_green(i)=S_green(i).Perimeter;
        N=N+1;
    end;
end;

%% Image proporties of the blue channel
N=1; nuclei_num=0;
for i=1:size(S_blue,1)
    if S_blue(i).Area>10
        orient_blue(i)=S_blue(i).Orientation;
        Eccentricity_blue(i)=S_blue(i).Eccentricity;
        complexity_blue(i)=1/(S_blue(i).Area/(S_blue(i).Perimeter^2)*4*pi);
         length_blue(i)=S_blue(i).MajorAxisLength;
         Perimeter_blue(i)=S_blue(i).Perimeter;
          Blue_size(i)=S_blue(i).Area; %nucleus size
          nuclei_num=nuclei_num+1; %counting number of nuclei
        N=N+1;
    end;
end;

%% average caculation for all parameters and channels
avg_blue_size=mean(Blue_size);

density_red=sum(sum(imBW_red))/size(im,1);
density_green=sum(sum(imBW_green))/size(im,1);
density_blue=sum(sum(imBW_blue))/size(im,1);
density_green_red=sum(sum(imBW_green_red))/size(im,1);
Nor_density_red=density_red/density_blue;
Nor_density_green=density_green/density_blue;


% Eccentricty analysis
Eccentricity0_25_red=100*sum((Eccentricity_red>0)&(Eccentricity_red<0.25))/size(Eccentricity_red,2); %this value will be of the most round elements
Eccentricity25_50_red=100*sum((Eccentricity_red>0.25)&(Eccentricity_red<0.5))/size(Eccentricity_red,2);
Eccentricity50_75_red=100*sum((Eccentricity_red>0.5)&(Eccentricity_red<0.75))/size(Eccentricity_red,2);
Eccentricity75_80_red=100*sum((Eccentricity_red>0.75)&(Eccentricity_red<0.8))/size(Eccentricity_red,2);
Eccentricity80_85_red=100*sum((Eccentricity_red>0.8)&(Eccentricity_red<0.85))/size(Eccentricity_red,2);
Eccentricity85_90_red=100*sum((Eccentricity_red>0.85)&(Eccentricity_red<0.9))/size(Eccentricity_red,2);
Eccentricity90_95_red=100*sum((Eccentricity_red>0.9)&(Eccentricity_red<0.95))/size(Eccentricity_red,2);
Eccentricity95_100_red=100*sum((Eccentricity_red>0.95)&(Eccentricity_red<1))/size(Eccentricity_red,2); %this value will be of the most elongated elements

%other channels
Eccentricity95_100_green=100*sum((Eccentricity_green>0.95)&(Eccentricity_green<1))/size(Eccentricity_green,2); %this value will be of the most elongated elements
Eccentricity95_100_blue=100*sum((Eccentricity_blue>0.95)&(Eccentricity_blue<1))/size(Eccentricity_blue,2); %this value will be of the most elongated elements

%orientation analysis for red
elements0_red=sum(abs(orient_red)<30);
elements90_red=sum(abs(orient_red)>60);
elementsOther_red=size(orient_red,2)-elements90_red-elements0_red;

%orientation variance
VarOri_red=var(abs(orient_red));

%orientation analysis for green
elements0_green=sum(abs(orient_green)<30);
elements90_green=sum(abs(orient_green)>60);
elementsOther_green=size(orient_green,2)-elements90_green-elements0_green;

%orientation variance
VarOri_green=var(abs(orient_green));

%orientation analysis for blue
elements0_blue=sum(abs(orient_blue)<30);
elements90_blue=sum(abs(orient_blue)>60);
elementsOther_blue=size(orient_blue,2)-elements90_blue-elements0_blue;

%orientation variance
VarOri_blue=var(abs(orient_blue));


%% Writing the results into a csv file
fid = fopen('results3.csv', 'a+');%name of excel file
fprintf(fid,'file name,Averaged nuclei size(pix),Normalized Red channel density,Normalized green channel density,Elements of eccentricity 0.95-1 in red,Elements of eccentricity 0.95-1 in green,Elements of eccentricity 0.95-1 in blue,Orientation variance in red,90 deg elements in red,Orientation variance in green,90 deg elements in green,Orientation variance in blue,90 deg elements in blue  \n');

fprintf(fid,'%s, %f, %f, %f, %f, %f, %f, %f, %f, %f, %f,%f, %f, \n',filename,avg_blue_size,Nor_density_red,Nor_density_green,Eccentricity95_100_red,Eccentricity95_100_green,Eccentricity95_100_blue,VarOri_red,elements90_red,VarOri_green,elements90_green,VarOri_blue,elements90_blue);
fclose(fid);
```
